# Toward closing a heartwater monitoring gap with a new pan-species ELISA suitable for domestic ruminants, wildlife, and camels

**DOI:** 10.64898/2026.10.29.762616

**Authors:** Valérie Rodrigues, Laetitia Viry, Alexis Monteil, Mélanie Dhune, Maelle Bayet, Philippe Pourquier, Valentin Ollivier, Fabien Donnet, Darshana Morar-Leather, Zinathi Dlamkile, Luis Neves, Damien F. Meyer

## Abstract

Heartwater, caused by the tick-borne bacterium *Ehrlichia ruminantium*, threatens ruminant production in sub-Saharan Africa and the Caribbean. Serological surveillance relies on species-specific ELISAs, so wildlife and camelids, which are potential reservoirs or sentinels, have no validated tools. We developed and evaluated a pan-species indirect ELISA prototype (ID Screen® *Ehrlichia ruminantium* Indirect Multi-species, Innovative Diagnostics) that detects anti-MAP1-B antibodies in all mammals. A positivity threshold of 30% relative positivity percentage (RPP) was empirically established from 707 heartwater-negative cattle, goat and sheep sera from three regions (maximum RPP 22.1%). Against a goat-specific reference ELISA (n = 503), the prototype showed 98.2% sensitivity, 97.2% specificity and 97.6% overall agreement. ROC analysis identified a Youden-optimal cut-off of 29.96%, confirming the empirical threshold. Inter-assay reproducibility was satisfactory (mean CV 7.4%), and qualitative classification remained stable across runs, including for a near-threshold sample. In four vaccinated sheep, the prototype detected seroconversion at the same time points as a sheep-specific ELISA. In non-domestic hosts, the assay gave uniformly low RPP values in 24 zoo animals of ten species from a heartwater-free area. By contrast, 4.5% of 111 dromedary camels from Chad exceeded the threshold. Eight wildlife species from Kruger National Park (n = 405) were tested and five species presented high median RPP values. This single-protocol assay provides a reproducible tool for integrated heartwater surveillance across livestock, camelids and wildlife, in Africa, in the Caribbean and in all countries threatened by the emergence of heartwater.

**Highlights:**

- A pan-species ELISA detects anti-MAP1-B antibodies across mammalian hosts
- Developed as a standardized commercial-format prototype (ID Screen® *Ehrlichia ruminantium* Indirect Multi-species ELISA, Innovative Diagnostics)
- 98.2% sensitivity and 97.2% specificity versus a species-specific ELISA
- Assay is reproducible (CV 7.4%) and allows tracking vaccine-induced seroconversion
- Assay detects *E. ruminantium* exposure in Kruger wildlife and Chadian camels

## Introduction

Heartwater, caused by the intracellular bacterium *Ehrlichia ruminantium* and transmitted by ticks of the genus *Amblyomma*, remains one of the major sanitary constraints on ruminant farming in sub-Saharan Africa and the Caribbean, generating substantial economic losses and trade restrictions on live animal movements (Allsopp, 2015; WOAH, 2025). Beyond its historical enzootic range, the disease continues to evolve: the emergence of new competent vectors, such as the invasive tick *Rhipicephalus microplus* progressively replacing *Amblyomma variegatum* in several West African countries, together with intensified livestock trade and climate change, is increasing the risk of pathogen introduction into previously disease-free areas (Biguezoton et al., 2016; Some et al., 2023). In the Caribbean, the recent detection of the disease in Les Saintes, an archipelago previously considered free of heartwater, provides a concrete illustration of the risk of inter-island spread (Dufleit et al., 2025). Heartwater could also concern other territories where competent ticks are present and where local breeds are susceptible, notably in the other Caribbean islands and in the American mainland (Dufleit et al., 2026, 2025; Kasari et al., 2010; Mahan et al., 2000; Nair et al., 2021)

Serological surveillance, alongside molecular detection, are crucial diagnostic approaches, and remains to date the only available tools able to reveal silent pathogen circulation before the occurrence of clinical cases, particularly in sentinel animals or in areas with an uncertain status WOAH, 2025). In-house ELISA assays, based on the recombinant MAP1-B protein, as iELISA (Mondry et al., 1998; Van Vliet et al., 1996), or cELISA (Bell-Sakyi et al., 2003; Jongejan et al., 1991; Katz et al., 1997) have shown satisfactory performance in domestic ruminants, with sensitivities and specificities generally ranging between 85 and 96% (Faburay et al., 2007). These tests nevertheless have important limitations for large-scale use: in-house ELISAs suffer from a lack of inter-laboratory standardization, and have been developed and validated on a species-by-species basis (cattle, sheep, or goats), each requiring a dedicated species-specific conjugate, which in practice means as many reagents and protocols as species to be tested.

This species-specific approach, while operational for domestic livestock, rapidly reaches its limits when surveillance needs to be extended to other hosts. A growing body of evidence highlights the potential role of wildlife and camelids as reservoirs or sentinels of *E. ruminantium*. African buffalo and several antelope species have long been recognized as competent hosts of the pathogen (Andrew and Norval, 1989; Peter et al., 1998, 2002), while serological studies conducted in the Sahel and the Horn of Africa have demonstrated exposure of dromedary camels to *E. ruminantium*, in some cases in association with poorly characterized clinical syndromes such as the so-called Acute Camel Death Syndrome, whose etiology has not yet been formally confirmed (Collins et al., 2022; Younan et al., 2021). Yet none of these non-domestic hosts currently benefits from a validated serological test: existing kits, calibrated with an anti-domestic-species conjugate, cannot be directly transposed to species as diverse as wild bovids, rhinoceroses, or camelids, effectively leaving these populations outside the scope of standardized serological surveillance, despite their potentially central role in the epidemiological cycle.

This critical shortcoming, which was highlighted by heartwater experts during the Gap Analysis workshop organized by the USDA-ARS in 2018 (Heartwater Report 2018 GAP analysis) is particularly problematic given that heartwater is increasingly recognized as an interspecies disease. In ecosystems where domestic ruminants, including camelids and wildlife, share grazing land and water points, surveillance restricted to livestock alone risks overlooking wild reservoirs that sustain pathogen circulation, thereby underestimating the true risk of disease emergence or re-emergence (Nair et al., 2021; Dufleit et al., 2026).

Developing a pan-species test, an indirect ELISA using a detection conjugate able to recognize immunoglobulins from a broad range of mammalian species, is therefore a direct methodological response to this need. Such a tool would allow, with a single protocol and a single positivity threshold, direct comparison of results obtained in cattle and small ruminants, and detect exposure of camelids, and multiple wildlife species for heartwater, thereby facilitating comparative seroprevalence studies and the integration of wildlife and camelids as sentinels in regional surveillance programs (WOAH, 2018). The aim of developing this new test is to address diagnostic gaps in wildlife surveillance, camelid screening, and the investigation of novel hosts - contexts in which no validated tool currently exists and developing a species-specific conjugate for every relevant species would be impractical or economically unfeasible.

It is in this context that a prototype of a pan-species indirect ELISA (ID Screen® *Ehrlichia ruminantium* Indirect Multi-species ELISA) was developed, in collaboration between Cirad (WOAH reference laboratory for heartwater, Guadeloupe, France) and the company Innovative Diagnostics (Grabels, France), targeting antibodies directed against the MAP1-B specific protein of *E. ruminantium*, the most specific antigen, common to all reference serological tests for this disease (Jongejan et al., 1993; Van Vliet et al., 1995, 1996). This work first aimed to establish a robust positivity threshold for domestic ruminants using a large collection of confirmed negative sera. It also evaluated the diagnostic performance of the prototype against a species-specific reference ELISA, as well as its repeatability and ability to track post-vaccination seroconversion kinetics. In a second step, the prototype was applied to wildlife samples from the SANParks Biobank in Skukuza, Kruger National Park, South Africa, field-collected camelid samples from Chad and French zoo animals maintained in a heartwater-free environment. These samples were used to assess the prototypes’ ability to discriminate exposed from unexposed populations. Overall, the results aim to demonstrate the value of a pan-species approach for integrated heartwater serological surveillance at the interface of livestock farming, pastoralism, and wildlife conservation.

## Materials and Methods

### Ethical considerations

Use of personal sample collections was authorized by the relevant institutional and national bodies, including ethical approval, material transfer agreements and/or export/import authorizations obtained from Montpellier Zoo, the LTA Hérault (France), and the competent authorities in Chad and South Africa.

### Serum samples

#### Domestic ruminants

##### Negative panel and threshold determination

A total of 707 serum samples from domestic ruminants (cattle, goats and sheep) with a confirmed heartwater-negative status were used to characterize assay specificity and to establish a positivity threshold. These included sera from France and Switzerland (n=649, personal Innovative Diagnostics collection), from sheep raised in Kenya and confirmed seronegative by the reference ELISA prior to a vaccination trial (n=32, personal Cirad collection), and from goats and sheep housed in tick-free CIRAD and INRAe (National Research Institute for Agriculture, Food and Environment, Petit Bourg) animal facilities in Guadeloupe (n=26).

##### Comparison panel

A panel of 503 goat serum samples was used to compare the ID Screen® *Ehrlichia ruminantium* Indirect Multi-species prototype with the goat-specific reference ELISA. This panel combined heartwater-negative and positive sera collected from goat herds in Guadeloupe, FWI – France, and archived in the Cirad-managed MiVeC biobank, all previously classified as positive or negative by the goat-specific reference ELISA.

##### Vaccination kinetics

Antibody kinetics following vaccination were monitored in four sheep raised in Kenya (Sheep 68, 80, 92 and 93), after immunization with a four-strain inactivated, adjuvanted heartwater vaccine. All animals were confirmed seronegative by the sheep-specific reference ELISA prior to vaccination. Blood samples were collected at day 0 (day of vaccination), 7, 14, 18, 28, 35 and 42 post-vaccination (day 42 sample not available for Sheep 80), and tested in parallel with the ID Screen® *Ehrlichia ruminantium* Indirect Multi-species ELISA prototype and the sheep-specific reference ELISA.

##### Wildlife and camelid samples

Negative-control wildlife sera were obtained from a private sera collection of 24 individuals belonging to ten species, including rhinoceros (*Ceratotherium simum simum*, n=6), gazella (*Nanger dama mhorr*, n=6), vicuña (*Vicugna vicugna, n=4*), bharal (*Pseudois nayaur, n=2*), oryx (*Oryx dammah*, n=1), giraffe (*Giraffa camelopardalis rotschildi, n=1*) camel (*Camelus dromedarius, n=1*), *zebra* (*Equus zebra hartmannae, n=1),* addax (*Addax nasomaculatus*, n=1) and Cape eland (*Tragelaphus oryx,* n=1), housed at Montpellier Zoo (France), a facility located in a heartwater-free area and with no history of exposure.

Wildlife sera (n = 405) were obtained from the SANParks Biobank in Skukuza, Kruger National Park, South Africa. Samples were collected between 2023 and 2024, and tested at the laboratories of the Department of Veterinary Tropical Diseases of the University of Pretoria (South Africa). Animals were selected based on their presumed susceptibility to infection and their occurrence in an endemic area. The samples comprised of sera from African buffalos (*Syncerus caffer*, n = 60), black rhinoceros (*Diceros bicornis*, n=60), blue wildebeests (*Connochaetes taurinus*, n = 60), impalas (*Aepyceros melampus*, n = 60), white rhinoceros (*Ceratotherium simum simum*, n=58), greater kudus (*Tragelaphus strepsiceros*, n=55), Giraffes (*Giraffa camelopardalis giraffa,* n=40), and nyalas (*Tragelaphus angasii,* n=12).

Field camelid sera (n=111) were collected from dromedary camels (*Camelus dromedarius*) in Chad, in collaboration with the Smithsonian Institution in 2023, in a region where heartwater occurs sporadically and where animals are sedentary.

All of these samples were tested with the ID Screen® *Ehrlichia ruminantium* Indirect Multi-species ELISA prototype only, as no validated species-specific reference ELISA is available for these species; the positivity threshold established for domestic ruminants (RPP = 30%) was applied for interpretation and discussion of the results.

### Reference (species-specific) ELISA

Results obtained with the pan-species prototype ELISA were compared with in-house, species-specific indirect ELISAs developed by Cirad, based on the MAP1-B antigen using an HRP-conjugated anti-species (goat or sheep) immunoglobulin as described elsewhere (Van Vliet et al., 1995). A goat-specific ELISA was used as the reference assay for the diagnostic sensitivity/specificity comparison in domestic ruminants, and a sheep-specific ELISA was used as the reference assay for the vaccination kinetics study. Positivity thresholds for the mono-species assays had been established previously and were applied as reported (22% RPP for the goat-specific ELISA, threshold given by the WOAH reference laboratory).

### Commercial ELISA prototype

Sera were tested using a prototype indirect ELISA, the ID Screen® *Ehrlichia ruminantium* Indirect Multi-species ELISA (Innovative Diagnostics, Grabels, France), developed in collaboration with the Cirad reference laboratory for heartwater (UMR ASTRE, Guadeloupe, France). The assay was designed to detect antibodies directed against the recombinant MAP1-B protein of *E. ruminantium*, and was intended for use with individual serum samples from any susceptible mammalian species. Briefly, microplate wells were coated with recombinant MAP1-B protein. Serum samples, together with positive and negative kit controls, were diluted 1/20 and dispensed into the wells. After incubation, plates were washed and a horseradish peroxidase (HRP)-conjugated anti-mammalian-IgG conjugate was added to detect bound anti-MAP1-B antibodies. Following a further washing step, the reaction was revealed using a tetramethylbenzidine (TMB) substrate solution, and optical density (OD) was read at 450 nm. For each sample, results were expressed as a relative positivity percentage (RPP), calculated as: RPP = (OD_sample_ / mean OD_positive control_) × 100.

### Reproducibility assay

Inter-assay reproducibility of the ID Screen® *Ehrlichia ruminantium* Indirect Multi-species ELISA prototype was assessed using five goat serum samples selected to span a range of RPP values, from near the positivity threshold to strongly positive. Each sample was tested in ten independent runs (1 plate per day, by the same operator), and reproducibility was expressed as the mean RPP, standard deviation (SD) and coefficient of variation (CV%, = SD/mean × 100) for each animal, together with the consistency of the qualitative (positive/negative) classification across runs relative to the RPP = 30% threshold.

### Statistical analysis

Differences in RPP values between geographic groups of confirmed heartwater-negative domestic ruminants were assessed using the Kruskal-Wallis rank-sum test. The positivity threshold for the ID Screen® *Ehrlichia ruminantium* Indirect Multi-species ELISA prototype was defined empirically from the distribution of RPP values in this negative population (mean, SD and maximum observed value), with a conservative margin above the highest observed value. This proposed positivity threshold was further evaluated by ROC curve analysis, using the classification obtained with the goat-specific reference ELISA (RPP ≥ 22%) as the reference standard. The area under the curve (AUC) was calculated with its 95% CI, and the Youden index (J = sensitivity + specificity − 1) was used to identify the statistically optimal cut-off, which was compared with the empirically retained threshold of RPP = 30%.

The association between RPP values obtained with the two assays was quantified using Pearson’s correlation coefficient (r, with 95% CI computed by Fisher’s z-transformation) and, because RPP values were not normally distributed (Shapiro-Wilk, p < 0.001), Spearman’s rank correlation coefficient (ρ). As correlation does not assess agreement, agreement between the two methods was evaluated by Passing-Bablok regression, an approach appropriate for comparing two methods that are both subject to measurement error, providing the slope and intercept with their 95% confidence intervals (CI). Diagnostic performance of the prototype assay relative to the goat-specific reference ELISA was summarized as a 2 × 2 contingency table, from which diagnostic sensitivity, specificity, positive and negative predictive values, and overall accuracy were calculated, each with exact (Clopper-Pearson) 95% CI.

All statistical analyses were performed using R statistical software. A p-value < 0.05 was considered statistically significant.

## Results

### Domestic ruminants

#### Threshold

A total of 707 serum samples from confirmed heartwater-negative domestic ruminants-cattle, goats, and sheep-were tested using the ID Screen® *Ehrlichia ruminantium* Indirect Multi-species ELISA prototype. The samples came from France/Switzerland (n = 649), Kenya (n = 32), and Guadeloupe (n = 26). The aim was to assess the specificity of the assay and establish an appropriate threshold for a positive result (Fig. 1). Relative positivity percentage (RPP) values ranged from 2.4% to 22.1%, with an overall mean of 6.6% (SD = 3.0%) and a median of 5.9%. Because none of the samples had a RPP above 22.1%, all 707 samples were classified as negative when a proposed threshold of 30% was applied. RPP values differed significantly between geographic groups (Kruskal-Wallis, H = 184.4, p < 0.001). Samples from France/Switzerland had the lowest median RPP value (5.6%; SD=2.8%), compared with samples from Kenya (10.7%; SD=1.5%) and Guadeloupe (10.9%; SD=2.1%). Nevertheless, all samples from all three regions remained well below the proposed 30% threshold. This indicates that the threshold is robust across different animal species and geographic locations. Based on these findings, RPP values of 30% was proposed as the positivity threshold. This value is approximately 7.8 standard deviations above the overall mean of the negative samples and provides a safety margin of 8 percentage points above the highest RPP value observed in confirmed negative samples.

**Fig. 1.**
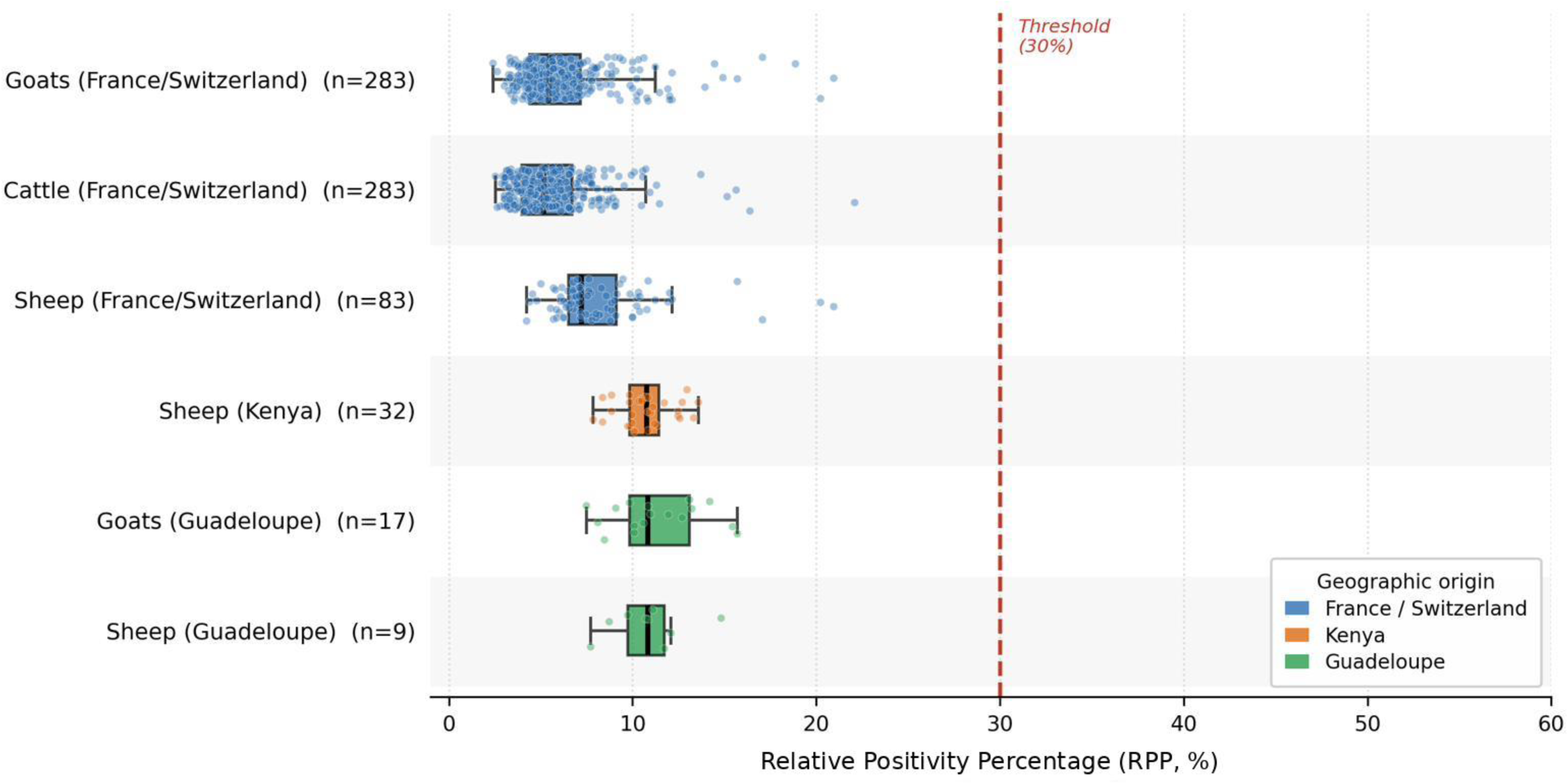
Distribution of Relative Positivity Percentage (RPP) values in heartwater-negative serum samples from domestic ruminants, tested with the ID Screen® Ehrlichia ruminantium Indirect Multi-species ELISA *prototype (n = 707)*. RPP = (OD*sample* / mean OD*pos.ctrl*) × 100. Samples were collected from cattle, goats, and sheep originating from heartwater-free areas in France/Switzerland (n = 649), Kenya (n = 32), and Guadeloupe (n = 26). Colors indicate geographic origin: blue, France/Switzerland; orange, Kenya; green, Guadeloupe. Groups are ordered by decreasing sample size. The box represents the interquartile range (IQR), the vertical bar the median, and whiskers extend to 1.5×IQR. All individual data points are shown. The dashed red line indicates the positivity threshold (RPP = 30%).

#### ELISA pan species vs specific ELISA comparison

A strong, highly significant positive correlation was observed between the RPP values obtained by mono-species and ID Screen® *Ehrlichia ruminantium* Indirect Multi-species ELISA (Pearson’s r = 0.905, 95% CI: 0.888–0.919, p < 0.001; n = 503) (Figure 2a) showing that both assays ranked samples consistently. This high overall correlation largely reflects the clear separation of negative and positive samples by both assays. Because correlation measures association rather than agreement, agreement between the two methods was further assessed by Passing-Bablok regression, an approach suited to comparing two assays each subject to measurement error, estimated a slope of 1.10 (95% CI: 1.04–1.16) and an intercept of −2.18 (95% CI: −2.75 to −1.67). As the slope was close to 1 and the intercept close to 0, these results suggest good overall agreement between the two methods, with no marked proportional or systematic bias, although a slight tendency for the pan-species assay to yield higher values than the mono-species assay was observed among strongly positive samples. Applying the positivity cut-offs defined for each assay (22% for mono-species and 30% for pan-species), 215 samples (42.7%) were classified as positive by both methods and 276 (54.9%) as negative by both methods, corresponding to an overall agreement of 97.6%. Only 12 samples (2.4%) yielded discordant results: 4 (0.8%) positive by the mono-species assay only and 8 (1.6%) positive by the pan-species assay only. These findings indicate good diagnostic agreement between the two ELISA formats.

**Fig. 2a.**
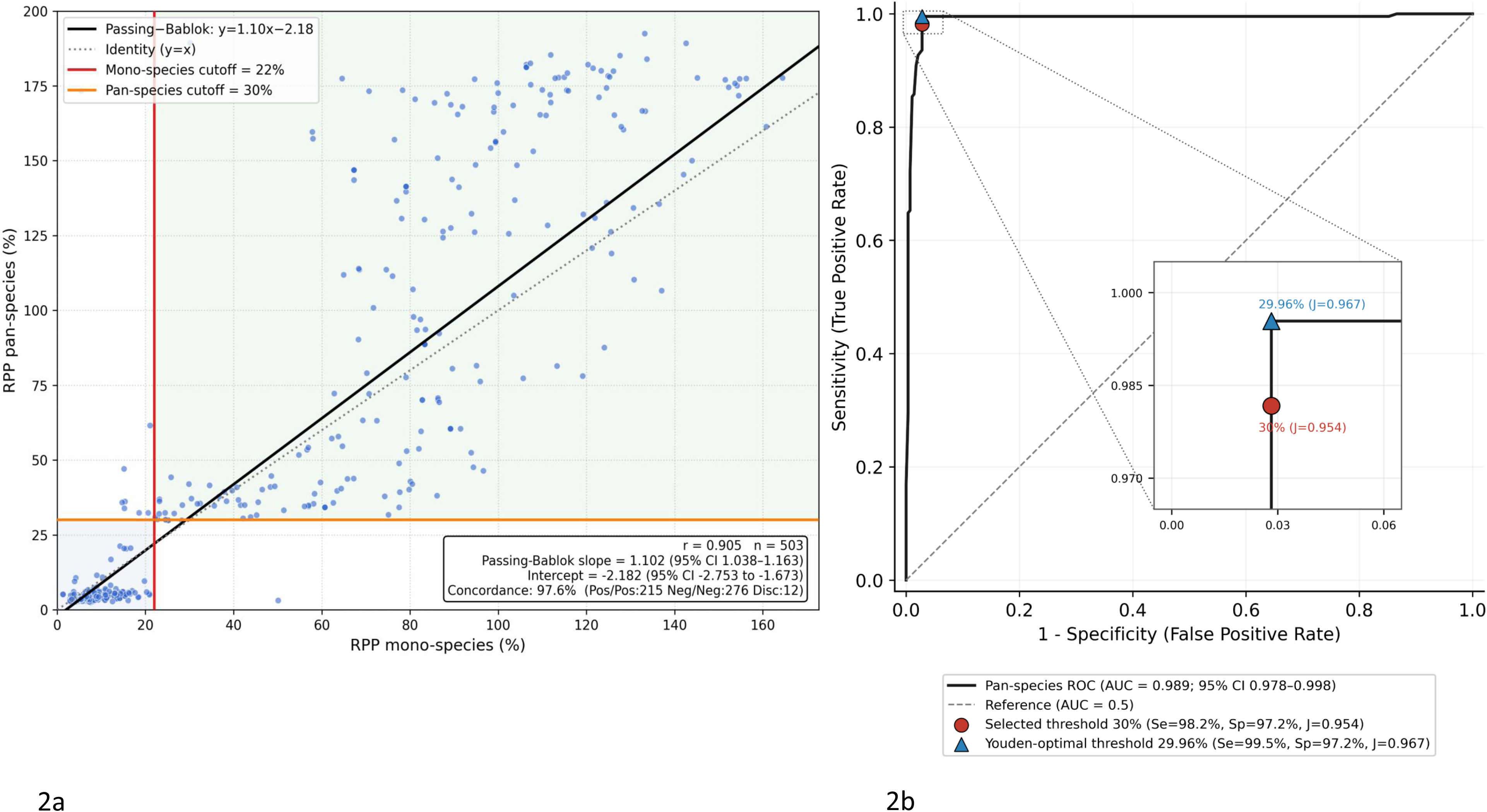
Correlation between the relative positivity percentage (RPP) values obtained by mono-species and ID Screen® Ehrlichia ruminantium Indirect Multi-species ELISA prototype for the detection of anti-MAP1-B antibodies (n = 503). Each point represents an individual serum sample. The solid black line represents the Passing-Bablok regression (y = 1.10x − 2.18; 95% CI of the slope: 1.04–1.16). The dotted grey line represents the line of identity (y = x), provided as a reference for assessing agreement between the two methods. The vertical (red) and horizontal (orange) lines indicate the positivity cut-offs used for the mono-species (22%) and pan-species (30%) assays, respectively, delineating four result zones: positive/positive (green background), negative/negative (blue background), and two zones of discordant results (white background) Fig. 2b. ROC curve for the ID Screen® Ehrlichia ruminantium Indirect Multi-species ELISA prototype, using the mono-species ELISA classification (RPP ≥ 22%) as reference (n = 503). AUC = 0.989 (95% CI: 0.978–0.998). Orange dot: currently retained threshold (RPP = 30%; Se = 98.2%, Sp = 97.2%, Youden J = 0.954). Green diamond: Youden-optimal threshold (RPP = 29.96%; Se = 99.5%, Sp = 97.2%, Youden J = 0.967). Insert: zoom on the upper-left region of the curve. The dashed diagonal line represents the line of no discrimination (AUC = 0.5).

#### Diagnostic performance

The diagnostic performance of the ID Screen® *Ehrlichia ruminantium* Indirect Multi-species ELISA prototype was evaluated by comparison with the goat-specific reference ELISA on a panel of 503 serum samples (Table 1). Of the 219 samples classified as positive by the reference assay, the ID Screen® *Ehrlichia ruminantium* Indirect Multi-species ELISA correctly identified 215 as positive (true positives) and incorrectly classified four as negative (false negatives). Among the 284 samples classified as negative by the reference assay, 276 were correctly identified as negative (true negatives), while 8 were incorrectly classified as positive (false positives). The diagnostic sensitivity of the ID Screen® *Ehrlichia ruminantium* Indirect Multi-species ELISA prototype, defined as its ability to correctly identify samples positive by the reference assay, was Se 98.2% (215/219; 95% CI: 95.4–99.5%). Its diagnostic specificity, defined as its ability to correctly identify samples negative by the reference assay, was 97.2% (276/284; 95% CI: 94.5–98.8%). The overall accuracy of the assay was 97.6% (95% CI: 95.9–98.8%). These results indicate an excellent level of agreement between the two assays, supporting the use of the ID Screen® *Ehrlichia ruminantium* Indirect Multi-species ELISA as a reliable alternative to species-specific serological tests for the detection of *E. ruminantium* MAP1-B antibodies in domestic ruminants.

**Table 1.**
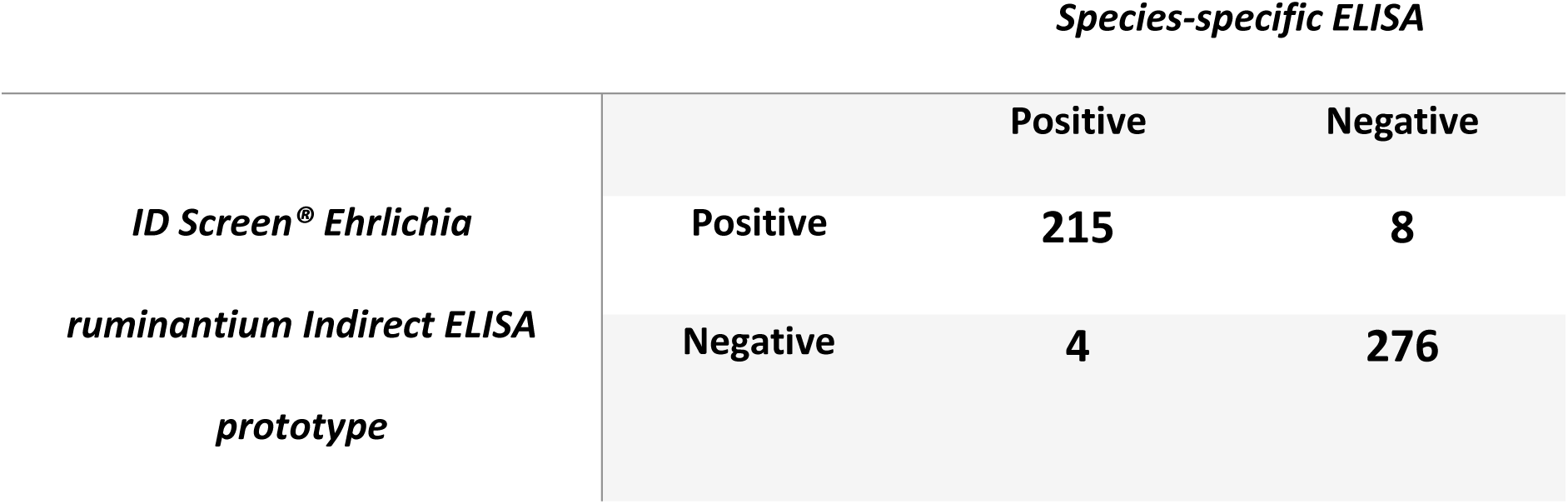
Comparative results of the ID Screen® Ehrlichia ruminantium Indirect Multi-species ELISA prototype and the mono-species goat-specific ELISA, on 503 serum samples. Diagnostic sensitivity and specificity of the ID Screen® Ehrlichia ruminantium Indirect Multi-species ELISA, calculated against the goat-specific ELISA as reference, were 98.2% (215/219; 95% CI: 95.4–99.5%) and 97.2% (276/284; 95% CI: 94.5–98.8%), respectively.

#### Threshold confirmation by ROC analysis

To further evaluate the proposed RPP positivity threshold of 30%, a ROC analysis was performed using the same panel of 503 samples used for the mono-species/pan-species comparison. Results from the mono-species ELISA, using a RPP cut-off of ≥ 22%, were used as the reference standard. The pan-species assay showed excellent discriminatory performance, with an area under the ROC curve (AUC) of 0.989 (95% CI: 0.978–0.998) (Fig 2b). The Youden index (J) was highest at a RPP threshold of 29.96% (J=0.967), corresponding to a sensitivity of 99.5% and a specificity of 97.2%. This cut-off was virtually identical to the empirically-derived threshold of RPP = 30% retained for this assay: at RPP = 30%: sensitivity = 98.2%, specificity = 97.2%, and J = 0.954.

#### Reproducibility on positive samples

The inter-assay reproducibility of the ID Screen® *Ehrlichia ruminantium* Indirect Multi-species ELISA prototype was assessed by testing five serum samples with contrasting RPP levels in 10 independent runs (Fig. 3). Mean RPP values ranged from 30.7% to 103.6% across the five goats, covering the full range from near-threshold to strongly positive results. Coefficients of variation (CV%) ranged from 0.0% to 14.2%, with a mean CV of 7.4% across all five goats. The lowest variability was observed for goat 6, which returned a constant RPP of 100% across all 10 runs (CV = 0.0%), and goat 13, which showed a CV of 7.9% despite having a mean RPP of 30.7%, immediately above the positivity threshold. Importantly, the qualitative result, positive or negative relative to the 30% threshold, remained consistent across all 10 runs for all five goats, with no run producing a discordant classification. The highest CV was observed for goat 1 (CV = 14.2%, mean RPP = 69.4%), which nonetheless remained unambiguously positive throughout.

**Fig. 3.**
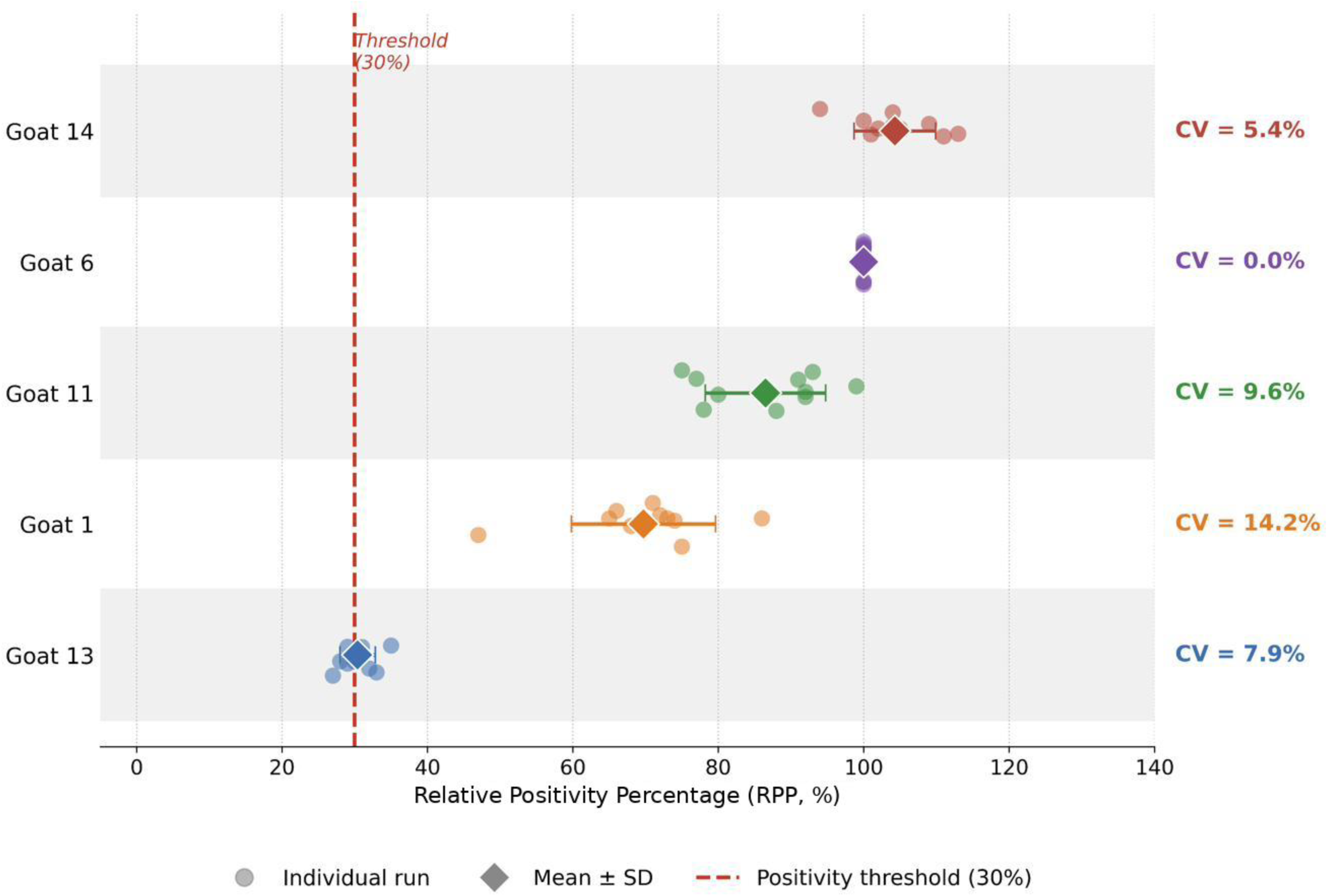
Inter-assay reproducibility of the ID Screen® Ehrlichia ruminantium Indirect Multi-species ELISA prototype evaluated on five goat serum samples tested in 10 independent runs. Each dot represents an individual run result. Diamonds indicate the mean RPP value for each animal; horizontal bars represent ± 1 standard deviation (SD). The coefficient of variation (CV%) is indicated for each animal. The dashed red line indicates the positivity threshold (RPP = 30%). RPP = (OD*sample* / mean OD*pos.ctrl*) × 100.

#### Antibody kinetics after vaccination

The antibody kinetics following vaccination against heartwater were monitored in four Kenyan sheep (Sheep 68, 80, 92, and 93) using both the ID Screen® *Ehrlichia ruminantium* Indirect Multi-species ELISA prototype and the sheep-specific reference ELISA, over a period of 42 days post-vaccination (Fig. 4). All animals tested negative with the sheep-specific reference ELISA before acclimatization at the facilities. Just prior to vaccination (D0), all four animals returned negative results with both assays, with RPP values well below the respective positivity thresholds of 30% (ID Screen® *Ehrlichia ruminantium* Indirect Multi-species ELISA) and 22% (sheep-specific ELISA), confirming their seronegative status at baseline. Seroconversion was detected by both assays in all four animals, with RPP values rapidly increasing from day 14–18 post-vaccination. Three animals (Sheep 80, 92, and 93) showed rapid and robust responses, ID Screen® *Ehrlichia ruminantium* Indirect Multi-species ELISA RPP values exceeding 80% by day 18 and reaching values of 141–158% from days 18-28 onward. Sheep 68 displayed a more moderate kinetic profile, crossing the ID Screen® *Ehrlichia ruminantium* Indirect Multi-species ELISA positivity threshold (30%) at day 18 and reaching a maximum RPP of 85% at day 42. Both ELISAs detected seroconversion at the same time points in all four animals, with RPP values exceeding the respective thresholds of 22% and 30% from days 14–18 onward. The ID Screen® *Ehrlichia ruminantium* Indirect Multi-species ELISA prototype generated higher absolute RPP values than the sheep-specific ELISA in all cases, consistent with differences in assay calibration, but the two tests crossed their respective thresholds at the same time point in the four animals. Sheep 80 showed a marked decline in RPP between days 28 and 42 with both assays, suggesting a more transient humoral response in this individual compared to the other three animals, in which RPP values remained elevated through day 42, and this decline was similarly detected by both ELISAs.

**Fig 4:**
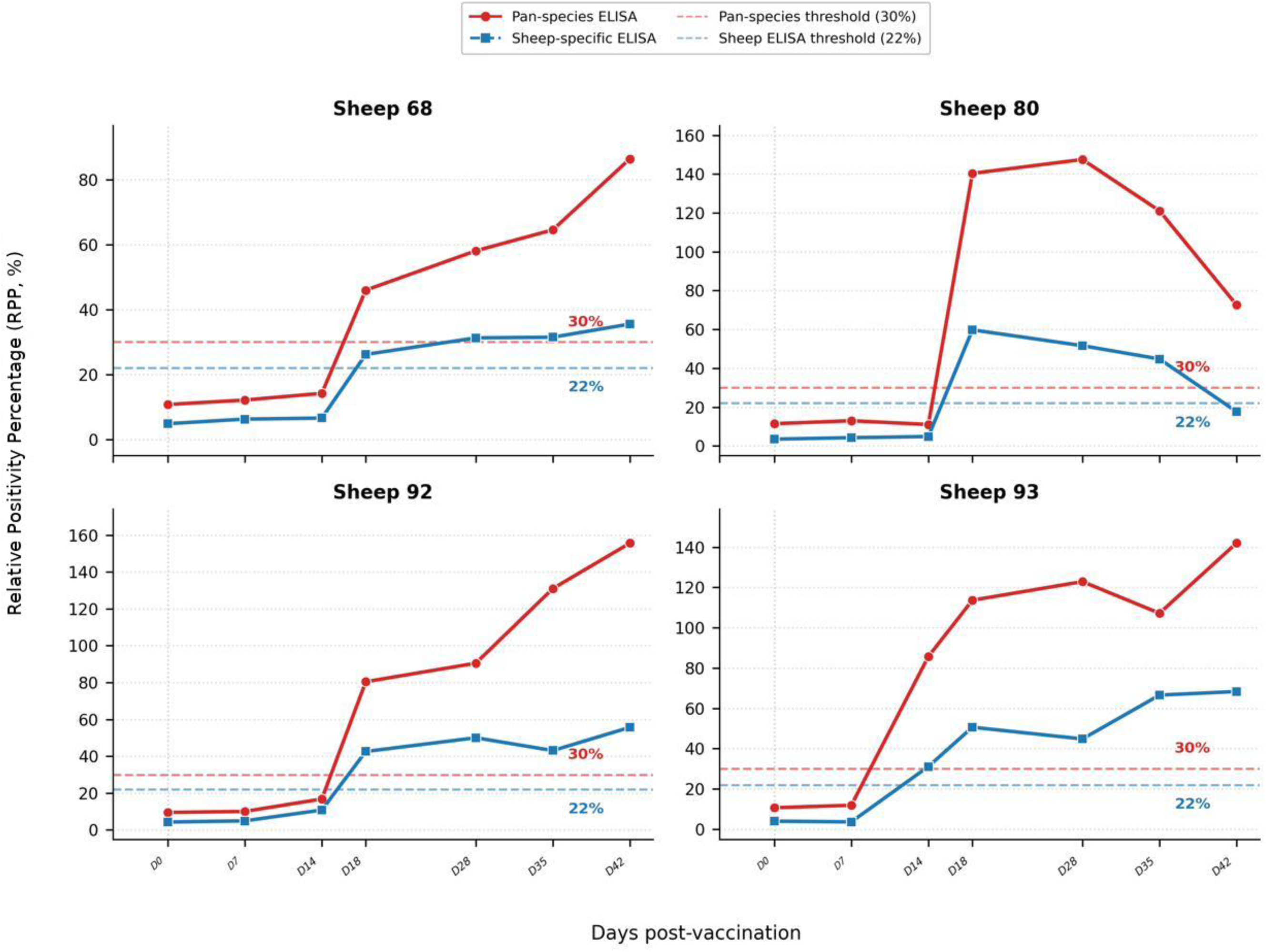
Antibody kinetics in four Kenyan sheep following vaccination against *Ehrlichia ruminantium*, detected by two serological assays. RPP (Relative Positivity Percentage = (OD*sample* / mean OD*pos.ctrl*) × 100) values are plotted over time for each animal individually. Red circles: ID Screen® *Ehrlichia ruminantium* Indirect Multi-species ELISA prototype; blue squares: sheep-specific ELISA. Dashed red line: positivity threshold for the ID Screen® *Ehrlichia ruminantium* Indirect Multi-species ELISA (RPP = 30%); dashed blue line: positivity threshold for the sheep-specific ELISA (RPP = 22%). D0 indicates the day of vaccination.

### Other mammals

#### Negative samples from heartwater-free area

Serum samples from 24 wildlife individuals, representing ten species, and housed in a heartwater free facility for many years, were tested with the ID Screen® *Ehrlichia ruminantium* Indirect Multi-species ELISA prototype. All animals returned RPP values ranging from 2.3% to 8.0% (median = 3.3%; Fig. 5), providing an overall estimate of the background for wildlife (n=24, 10 species).

**Fig. 5.**
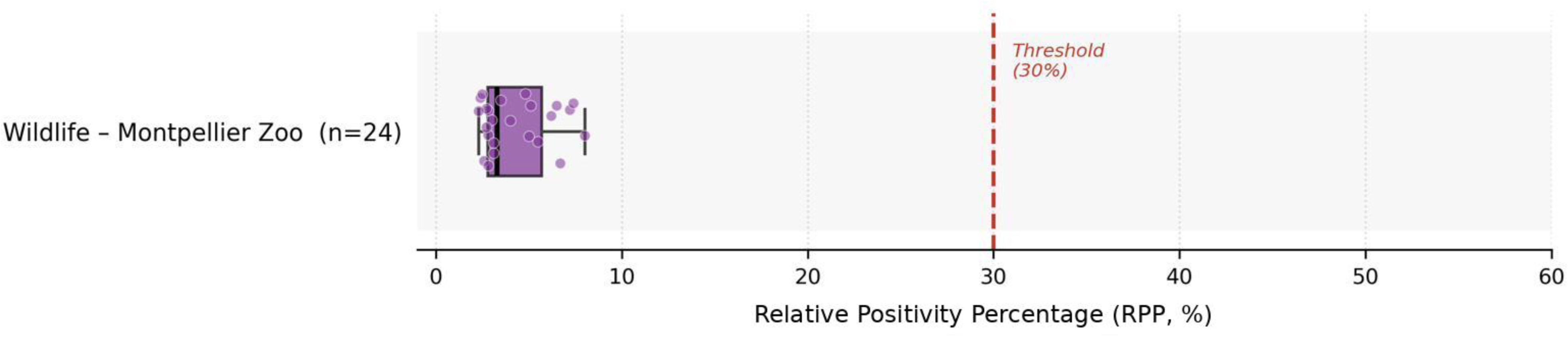
Distribution of Relative Positivity Percentage (RPP) values in heartwater-negative wildlife serum samples collected from the Montpellier Zoo (France), tested with the ID Screen® *Ehrlichia ruminantium* Indirect Multi-species ELISA prototype (*n* = 24). RPP = (OD*sample* / mean OD*pos.ctrl*) × 100. The box represents the interquartile range (IQR), the vertical bar the median, and whiskers extend to 1.5×IQR. Individual data points are shown as jittered dots. The dashed red line indicates the ruminants’ positivity threshold.

#### Field camelids

A total of 111 dromedary camel serum samples collected in Chad were tested with the ID Screen® *Ehrlichia ruminantium* Indirect Multi-species ELISA prototype. RPP values ranged from 2.3% to 72.8% (median = 6.1%; Fig. 6), with the majority of animals (106/111; 95.5%) giving results below the domestic ruminant-derived positivity threshold of 30%. However, five individuals (4.5%) showed RPP values of 36.8%, 39.0%, 40.8%, 48.4%, and 72.8%, respectively, which is higher than results obtained from the large majority of tested animals and from zoo-kept animals in a heartwater-free area, where no individual exceeded 8.0%. Nine animals, in addition to the previous 5, also had high RPP values (above the Q3 + 1.5 IQR); these results should be interpreted with caution, as they could indicate low seropositivity or cross-reaction with another pathogen.

**Fig. 6.**
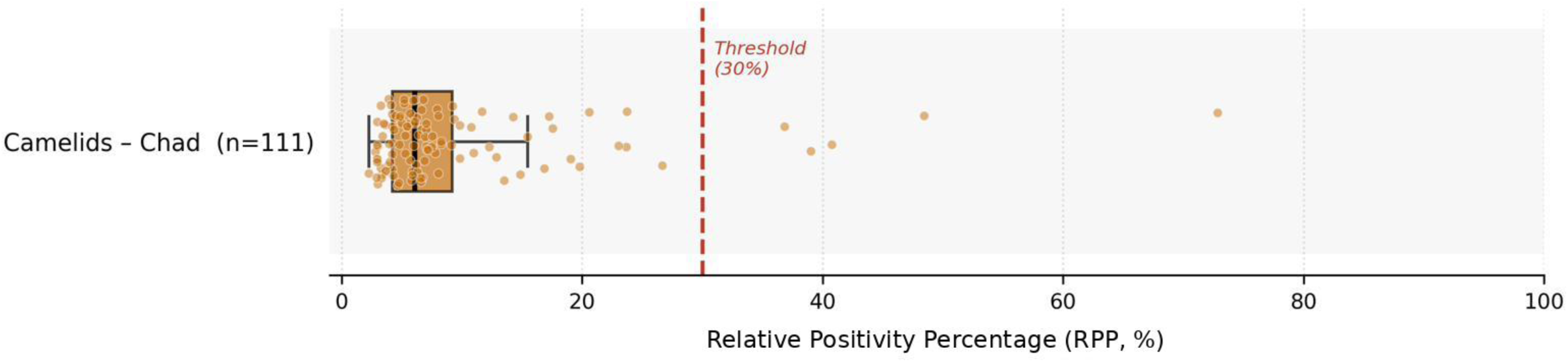
Distribution of Relative Positivity Percentage (RPP) values obtained with the ID Screen® *Ehrlichia ruminantium* Indirect Multi-species ELISA prototype in serum samples collected from dromedary camels (Camelus dromedarius) in Chad (*n* = 111). RPP = (OD*sample* / mean OD*pos.ctrl*) × 100. The box represents the interquartile range (IQR), the vertical bar the median, and whiskers extend to 1.5×IQR. Individual data points are shown as jittered dots. The dashed red line indicates the ruminants’ positivity threshold.

#### Wildlife species kept under natural conditions

The ID Screen® *Ehrlichia ruminantium* Indirect Multi-species ELISA was used to test 405 serum samples collected from eight wildlife species in Kruger National Park, South Africa. RPP values varied widely both within and among species (Fig. 7), ranging from near zero to 290.5%. Five species showed median RPP values above the domestic ruminant-derived threshold of 30%: white rhinoceros (103.4%, n = 58), nyala (90.8%, n = 12), greater kudu (64.5%, n = 55); giraffe (40.8%, n = 40) and African buffalo (32.5%, n = 60), in which about half of the individuals exceeded the cut-off. In contrast, blue wildebeest (median RPP = 14.6%, n = 60), black rhinoceros (14.5%, n = 60), and impala (9.7%, n = 60) showed median RPP values well below the threshold, although a small number of individuals in each group exceeded 30%. Given that these wildlife species were tested for the first time using this assay and that the biological significance of the results obtained cannot be linked to a possible infection, no statistical tests were applied to compare the results of the different species tested.

**Fig. 7.**
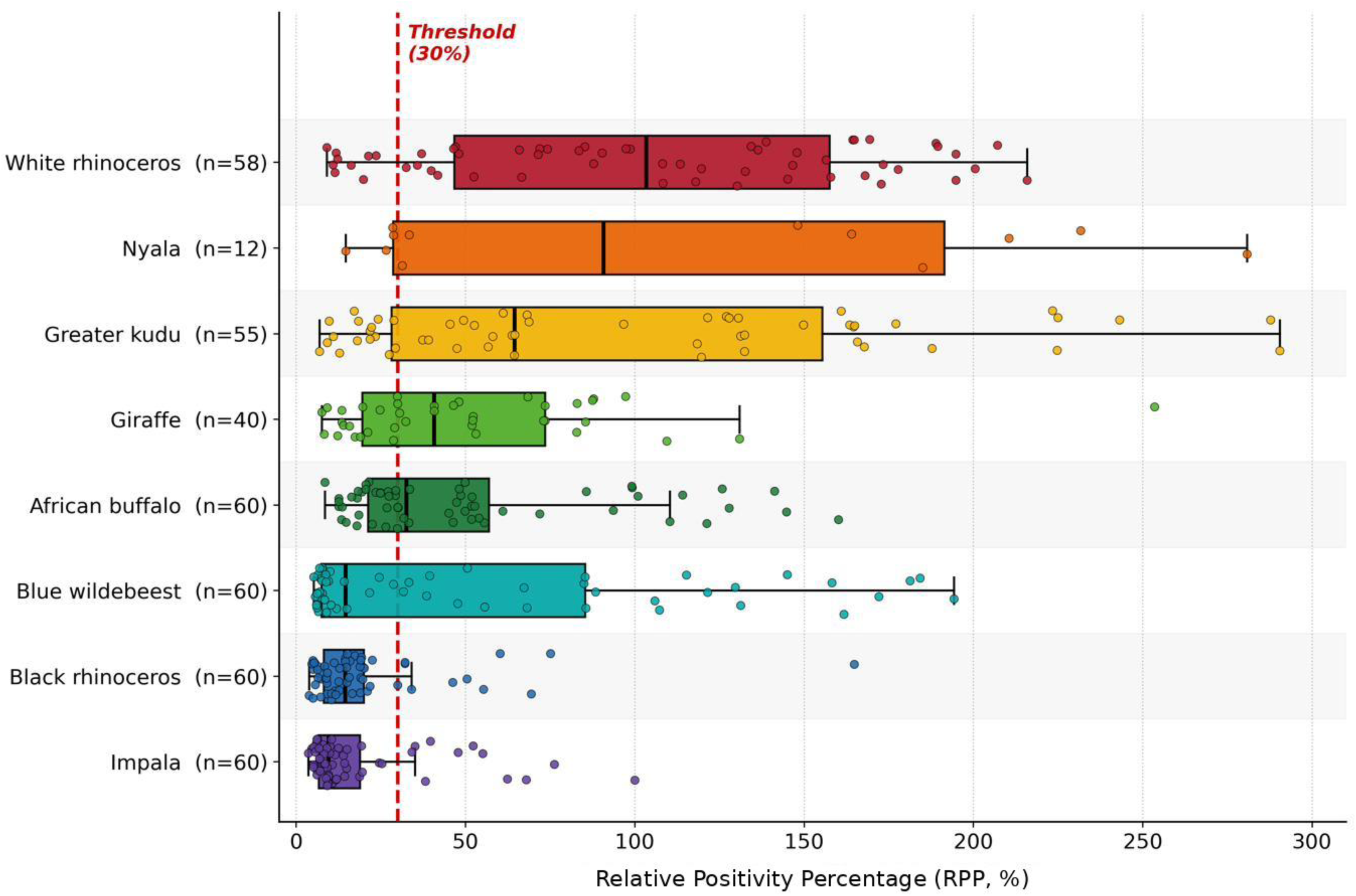
Distribution of Relative Positivity Percentage (RPP) values by Kruger Park (SA) wildlife species tested with the ID Screen® *Ehrlichia ruminantium* Indirect Multi-species ELISA prototype (*n* = 405). RPP = (OD*sample* / mean OD*pos.ctrl*) × 100. Species are ordered by decreasing median RPP. Boxes represent the interquartile range (IQR), vertical bars the median, and whiskers extend to 1.5×IQR. Individual data points are shown. The dashed red line indicates the ruminants’ positivity threshold.

## Discussion

This study provides the first evaluation of a pan-species indirect ELISA, developed in close collaboration between Cirad Guadeloupe, France, the WOAH reference laboratory for Heartwater and Innovative Diagnostics, Grabels, France, a company with advanced expertise in veterinary diagnostics, for the detection of anti-MAP1-B antibodies against *Ehrlichia ruminantium*, covering not only domestic ruminants but also wildlife and camelids. Our results demonstrate that this single assay combines strong performance in cattle, sheep, and goats with the ability to detect antibodies across a broad taxonomic range of hosts, addressing a long-standing gap in heartwater serological surveillance tools (Heartwater Report 2018 GAP analysis)

In domestic ruminants, the positivity threshold defined for this test, fixed at Relative Positivity Percentage (RPP) = 30%, proved highly robust, with all 707 confirmed heartwater-negative samples - from three geographically and epidemiologically distinct populations (France/Switzerland, Kenya, Guadeloupe) - remaining well below this cut-off (maximum RPP = 22.1%). This RPP threshold was confirmed by the ROC analysis with the Youden index, that was convergent with our first empirical approach: the distribution-based threshold derived from 707 confirmed heartwater-negative domestic ruminant sera (Fig. 1), and this ROC-optimal threshold derived from the 503-sample diagnostic comparison panel provides strong support for the robustness of the RPP = 30% cut-off for classification with the ID Screen® *Ehrlichia ruminantium* Indirect Multi-species ELISA prototype. Although the optimal cut-off determined by ROC analysis was set at 30% RPP, a positivity threshold of 50% RPP will be recommended in the kit instructions. This more conservative threshold was chosen to provide a safety margin that minimizes the risk of false-positive results, which is particularly important in heartwater-free areas, where a single false-positive result could have serious consequences for animal movements and trade. This choice is consistent with the intended use of the assay, which is designed for herd-level or area-level surveillance based on an appropriate sampling strategy rather than for individual diagnosis. In this context, two considerations support the use of the higher threshold. First, in an infected herd or area, sera with RPP values between 30% and 50% are very likely to be accompanied by other samples with strongly positive results, so the infection status of the population should still be correctly identified. Second, isolated sera with RPP values in this intermediate range may reflect either very early seroconversion or decrease in antibody levels at the end of infection. Such results should therefore not be disregarded: any suspicion of seropositivity should prompt the collection of *Amblyomma* ticks, such as *Amblyomma variegatum or Amblyomma hebraeum*, in the area concerned, followed by detection of *Ehrlichia ruminantium* DNA using the pCS20 quantitative PCR assay to confirm the circulation of the pathogen in the vector population (Cangi et al., 2017).

Although RPP values differed significantly between origins (Kruskal-Wallis, p < 0.001), with slightly higher background reactivity in animals from tropical settings (Kenya, Guadeloupe) compared with France/Switzerland, this variability did not compromise the specificity of the assay, since the threshold retained a margin of approximately 7.8 SD above the mean of the negative population. This finding is consistent with the general expectation that this background may be influenced by the different origins or breeds of animals or by non-specific low cross-reactivity, as previously described by using MAP1-B based ELISA (Kelly et al., 1994; Van Vliet et al., 1996; Mondry et al., 1998) but it confirms that a single, conservative threshold can be applied across diverse domestic ruminant populations without loss of specificity.

The diagnostic sensitivity (98.2%) and specificity (97.2%) obtained against the goat-specific reference ELISA are comparable to those reported for other MAP1-B-based serological assays, including an in-house ELISA, for example iELISA described by (Van Vliet et al., 1995; Mondry et al., 1998), and cELISA described by (Jongejan et al., 1991; Bell-Sakyi et al., 2003), which typically report sensitivities and specificities in the 85–96% range. The pan-species prototype demonstrated a strong 97.6% agreement with the species-specific reference assay, supported by narrow 95% confidence intervals. Since both assays use the same *E. ruminantium* antigen which is the most specific at this time, this validates the new test as a dependable alternative to species-specific tests, with the added operational advantage of a single protocol applicable across all domestic species. The 12 discordant results (2.4%) are noteworthy: the tendency of the pan-species assay to yield systematically higher RPP values than the specific assay (regression slope = 1.10) suggests that most discordances occurred in samples with borderline antibody titers close to either threshold, rather than reflecting a fundamental lack of analytical agreement. It should also be noted that species-specific results were generated several years prior to pan-species testing, and sample degradation over time cannot be excluded as a contributing factor to some of the observed false-negative results by the pan-species assay.

Inter-assay reproducibility was satisfactory, with a mean coefficient of variation of 7.4% across samples spanning the diagnostic range, which meets the criteria for reproducibility of results recommended in serological validation standards (ideally 10% - 15% max; Jacobson, 1998; WOAH, 2025) including one sample with a mean RPP value immediately above the positivity threshold. Critically, qualitative classification (positive/negative) remained stable across all ten replicate runs for all five animals tested, including this borderline sample, indicating that the assay’s classification is robust even in the zone of diagnostic uncertainty (*i.e.,* near-threshold results). This is an important practical consideration, since borderline results are precisely those most likely to affect surveillance decisions in the field, even though it is generally recommended that serology results for heartwater must be interpreted at the herd or regional level, rather than on an individual basis (WOAH, 2018). In fact, it has been shown, particularly in cattle, that serological prevalence remains lower than expected in endemic areas with repeated infections (Semu et al., 2001; Kakono et al., 2003; Simbi et al., 2003) and that it persists for a shorter period than small ruminants (Kakono et al., 2003; Faburay et al., 2005, 2007)

The antibody kinetics data obtained for this limited cohort of experimentally vaccinated sheep provide additional support for the assay’s diagnostic validity. Both the pan-species prototype and the sheep-specific ELISA detected seroconversion at concordant time points (day 14–18 post-vaccination) in all four animals, despite the pan-species assay consistently generating higher absolute RPP values, which could be attributed to differences in assay calibration. The moderate response observed in one animal (Sheep 68), together with the marked decline in titers observed between days 28 and 42 in another (Sheep 80), illustrate the individual variability of humoral responses following vaccination and underline the value of the pan-species ELISA as a tool capable of capturing this variability with a temporal resolution comparable to that of established species-specific assays.

Perhaps the most novel contribution of this study lies in the extension of ELISA testing capacity to non-domestic hosts. The contrast observed between wildlife individuals housed in a heartwater-free environment (Montpellier Zoo; RPP range 2.3–8.0%) and those sampled under field conditions in enzootic or potentially exposed areas (Chad camelids, Kruger National Park wildlife) provides indirect evidence that the assay is capturing signal related to *E. ruminantium* exposure, rather than non-specific reactivity of breeds other than domestic ruminants as camels or wildlife.

In dromedary camels sampled in Chad, a small proportion of animals (4.5%) exceeded the positivity threshold previously determined for domestic ruminants, with RPP values well above those observed in any zoo-housed, heartwater-free camelid. While these results should be interpreted with caution in the absence of molecular confirmation, they are consistent with previous serological evidence of *E. ruminantium* exposure in camelids from the Sahel and Horn of Africa, where clinical cases, named Acute Camel Death Syndrome, resembling heartwater, but in which the etiology has not yet been formally confirmed, (Younan et al., 2021; Collins et al., 2022) and are compatible with the hypothesis that camels may also act as under-recognized sentinels or possibly reservoirs of the pathogen in pastoral ecosystems where they share grazing land and water points with small ruminants.

The results obtained in free-ranging wildlife from the Kruger National Park were the most striking, revealing marked interspecies heterogeneity in results, from near-zero to extremely high RPP values (up to 290.5%). White rhinoceros, Nyala, and Greater kudu displayed high median RPP values several-fold above the positivity threshold applied to domestic ruminants, suggesting strong serological responses and/or frequent, and possibly chronic or repeated, exposure to *E. ruminantium* and its tick vectors in these species. Giraffe and African buffalo showed intermediate patterns, with approximately half of the buffalo samples surpassing the cut-off. This aligns with buffaloes’ established role as competent reservoir hosts (Andrew and Norval, 1989; Peter et al., 2002, 1998) and supports the suspected role of giraffes’ as long-term carrier of Ehrlichia ruminantium (Peter et al., 1998). In contrast, blue wildebeest, black rhinoceros, and impala showed lower median values, although a subset of individuals in each group exhibits high RPP values, indicating possible exposure even within a single ecosystem. This inter-species gradient raises an important methodological question: whether this single threshold, derived from domestic ruminant data, is appropriate for interpreting results across wildlife species, or whether species- or regional-specific cut-offs will ultimately be required, as proposed by others (Peter et al., 2002; Garnier et al., 2017). In the absence of confirmed negative and positive reference populations for most wildlife species, and without molecular confirmation of active infection, the wildlife results presented here should be regarded as indicative of species susceptibility and exposure at the population level, and, as for ruminant interpretation results, but certainly not as validated individual diagnostic outcomes.

Several limitations of this study should be acknowledged. First, because we used personal or bio banked collections of sera, seropositivity in wildlife and camelid samples could not be corroborated by PCR or pathogen isolation, precluding a definitive distinction between active infection, past exposure, and possible cross-reactivity with other *Ehrlichia* species that circulated in the region. Second, as previously mentioned, the positivity threshold was derived exclusively from domestic ruminant data and extrapolated to other species; although the zoo-derived negative samples support its applicability beyond domestic ruminants, sample sizes for most wildlife species remain limited (in some cases n = 1), constraining the robustness of species-specific results. Third, field samples from Chad and Kruger National Park were collected by partners for other scientific purposes, were selected based on suspected susceptibility in endemic areas, and were kindly made available for the validation of this test, which introduces a non-random sampling bias and must be taken into account in the epidemiological interpretation of the results.

Despite these limitations, the results of this study support the use of the ID Screen® *Ehrlichia ruminantium* Indirect Multi-species ELISA prototype as a valuable and versatile tool for heartwater epidemiological surveillance across domestic and non-domestic hosts. Its strong performances in domestic ruminants, combined with its capacity to discriminate exposed from unexposed individuals in wildlife and camelids, positions it as a candidate tool for integrated, cross-species surveillance strategies at the interfaces between livestock, wildlife, and pastoral systems, which is recognized as critical and essential for the heartwater epidemiology, regarding evolution in endemic areas or introduction into free areas. To confirm these serological observations, future work should include molecular confirmation of the pathogen *E. ruminantium*, either by tick sampling in the same areas or, for the definitive proof of infection, ideally when clinical signs are present, by sampling animals for molecular detection of *E. ruminantium*, even though it has already been shown that bacteremia fluctuates greatly during infection (Simbi et al., 2003).

This new simple, reproducible and reliable test, will make it possible to implement large-scale, integrated regional or international surveillance programs for heartwater, as future programs should systematically include local wildlife and/or camels as sentinels. Such an approach is critical for anticipating emergence risks and protecting livestock sectors in Africa, in the Caribbean and in all countries threatened by the emergence of heartwater.

## Acknowledgments

The authors are very grateful to all the partners for sharing samples for analysis: LTA Herault (Montpellier, France), Zoo de Montpellier (Montpellier France), breeders and technicians from animal facilities (Guadeloupe and Montpellier-France, Switzerland), Multivacc project (IDRC, Canada) for vaccination samples, and RFOARA project (Smithsonian Institution, USA) for camel samples.

We would like to extend our sincere thanks to the Innovative Diagnostics team (Grabels, France), which optimized the reagents for this test and produced the prototypes, using their industrial process to ensure their availability and reproducibility.

We would also like to thank Ms. Géraldine Bossard, from CIRAD - Intertryp research unit, for the informal discussions we had at the very beginning of this project, which convinced us to try this methodology, applicable to all species, for monitoring heartwater disease.

## Conflict of interest

The authors declare no competing interests.

The ID Screen® *Ehrlichia ruminantium* Indirect Multi-species ELISA prototype was co-developed by Cirad and Innovative Diagnostics, which supplied the prototype kits. Innovative Diagnostics had no role in study design, data analysis or interpretation. The corresponding author can provide the datasets for this work upon reasonable request.

## Declaration of generative AI and AI-assisted technologies

During the preparation of this work, the authors used Claude (Anthropic) to proofread the English and to assist in writing the R scripts used to plot figures from the original data. After using this tool, the authors reviewed and edited the content as needed and take full responsibility for the content of the publication.

